# Recirculation of H_2_, CO_2_, and ethylene improves carbon fixation and carboxylate yields in anaerobic fermentation

**DOI:** 10.1101/2021.06.11.448067

**Authors:** Flávio C. F. Baleeiro, Sabine Kleinsteuber, Heike Sträuber

## Abstract

Anaerobic fermentation with mixed cultures has gained momentum as a bioprocess for its promise to produce platform carboxylates from low-value biomass feedstocks. Anaerobic fermenters are net carbon emitters and their carboxylate yields are limited by electron donor availability. In a new approach to tackle these two disadvantages, we operated two bioreactors fed with acetate and lactate as a model feedstock while recirculating H_2_/CO_2_ to stimulate concomitant autotrophic activity. After 42 days of operation, hydrogenotrophic methanogenesis was predominant and ethylene (≥1.3 kPa) was added to one of the reactors, inhibiting methanogenesis completely and recovering net carbon fixation (0.20 g CO_2_ L^−1^ d^−1^). When methanogenesis was inhibited, exogenous H_2_ accounted for 17% of the consumed electron donors. Lactate-to-butyrate selectivity was 101% (88% in the control without ethylene) and lactate-to-caproate selectivity was 17% (2.3% in the control). Community analysis revealed that ethylene caused *Methanobacterium* to be washed out, giving room to acetogenic bacteria. In contrast to 2-bromoethanosulfonate, ethylene is a scalable methanogenesis inhibition strategy that did not collaterally block *i-*butyrate formation. By favoring the bacterial share of the community to become mixotrophic, the concept offers a way to simultaneously increase selectivity to medium-chain carboxylates and to develop a carbon-fixing chain elongation process.

## INTRODUCTION

Carboxylate production via anaerobic fermentation of complex biomass feedstocks, such as lignocellulose or food waste, recovers value-added products from low value waste streams^1^. Among the most common carboxylates produced, medium-chain carboxylates (MCC, e.g. caproate and caprylate) have received particular attention. MCC stand out as promising platform chemicals and sustainable energy carriers compared to short-chain carboxylates (SCC) and ethanol due to their higher energy density and low water solubility enabling an easier separation from fermentation broths^2–3^.

MCC production by mixed cultures results from the process of chain elongation in which linear carboxylate carbon chains are extended by two carbon atoms in each cycle. Chain elongation requires an electron donor and an electron acceptor (typically an SCC). Conventional electron donors (i.e. lactate, sugars, or ethanol) and SCC are produced by hydrolysis and fermentation of biomass^4^.

As a drawback, anaerobic fermenters are commonly net carbon emitters because some carbon from the feedstock is released in form of CO_2_ and CH_4_ via various metabolic pathways. CO_2_ is formed during fermentation of substrates such as carbohydrates and lactate into SCC via pyruvate (shown in Equation 1 for acetate); H_2_/CO_2_ is formed by syntrophic bacteria during SCC oxidation (shown in Equation 2 for acetate); CH_4_ and CO_2_ are produced from acetate by acetoclastic methanogens (Equation 3); and CH_4_ is formed from H_2_/CO_2_ by hydrogenotrophic methanogens (Equation 4).

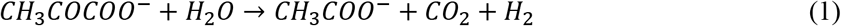

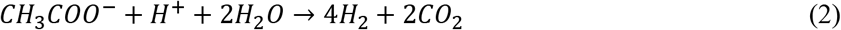

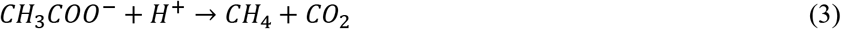

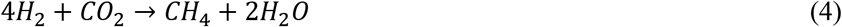

The pathways described by Equations 2, 3, and 4 are counterproductive to carboxylate production and chain elongation^5^ whereas CO_2_ emission in Equation 1 is a consequence of the stoichiometry in the production of carboxylates with even carbon numbers. If additional H_2_ is provided to the mixed culture, carbon emission can be compensated by homoacetogenic activity with reincorporation of CO_2_ into acetate (Equation 5).

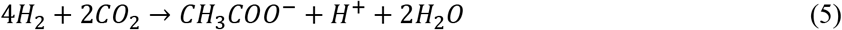

If H_2_ can be kept abundant and accessible to the microorganisms, multiple positive effects can be achieved by i) stimulating autotrophic activity in the community^6^ to the point that fermentation can become a net carbon-fixing process (with exogenous CO_2_); ii) disfavoring the oxidation of SCC (Equation 2)^5^; and iii) providing substrates for chain elongation. To take profit from these possibilities, the concept of an anaerobic fermenter with recirculation of exogenous H_2_/CO_2_ or syngas (H_2_, CO_2_, CO) was proposed Baleeiro, et al. ^5^.

To avoid carbon emission by methanogenesis (Equations 3 and 4), the use of chemical inhibitors is a popular alternative. Among methanogenesis inhibitors considered to be selective, 2-bromoethanesulfonate (2-BES) is the most used chemical in lab-scale fermentations^7^. However, the selectivity of 2-BES comes with caveats. Recent studies indicated collateral effects of 2-BES addition on carboxylate production, such as inhibition of *i-*butyrate formation^6, 8^ and an increase of sulfate-reducing bacterial populations, suggesting 2-BES degradation in the long term^9–11^. Moreover, application of 2-BES at the high concentrations (50 mM^12^) needed to inhibit hydrogenotrophic methanogenesis (Equation 4) might be economically unfeasible in industrial scale.

Alternatively, ethylene and acetylene are commodity gases that can inhibit methanogenesis completely even at partial pressures as low as 0.5 kPa^13^. Up to 5 kPa of ethylene showed no inhibitory effect on pure cultures of the acetogenic bacterium *Acetobacterium woodii*^14^, yet, studies on the use of these gases in anaerobic fermentation are rare, possibly because the gas phase of many reactors is simply vented out. In a few concept demonstrations, acetylene has been proposed as a cost-effective methanogenesis inhibitor in H_2_ production^15–17^. To the best of the author’s knowledge, there are no reports testing the cost-effectiveness of ethylene in the literature.

In this study, a gas recirculation system was developed with the aims of achieving net carbon fixation and enhancing carboxylate production with exogenous H_2_/CO_2_ without increasing the supply of conventional electron donors. To deal with the resilient methanogenic activity, the use of ethylene as a methanogenesis inhibitor was tested in culture bottles and scaled up to a H_2_/CO_2_ recirculation reactor.

## MATERIAL AND METHODS

### BATCH EXPERIMENTS IN CULTURE BOTTLES

Two different culture bottle experiments were realized with duplicates to test the use of ethylene as methanogenesis inhibitor. The first batch lasted 48 days with H_2_ (160 kPa) as electron donor and under conditions with and without ethylene. The second batch with addition of H_2_ (160 kPa) and ethanol (1.7 g L^−1^) as electron donors lasted 63 days, and conditions with ethylene, with 2-BES, and without methanogenesis inhibitor were tested. Table S1 summarizes the tested conditions and the types of controls used in each batch experiment.

The inoculum for the batch experiments originated from a previous study, in which microbial communities were enriched on organic substrates and H_2_/CO_2_^6^. From this study, “community C” was used for inoculation. Prior to their use, the inoculum sources were stored in serum bottles initially with 200 kPa H_2_:CO_2_ (80:20) in the dark and at room temperature. The basal medium containing yeast extract and 200 mM acetate was described by Baleeiro et al.^6^. The cultures were set up with 10 vol% inoculum, whereas the abiotic control bottles received 10 vol% sterile anoxic water instead. Preparation procedures for the fermentation and gas purging/pressurization cycles were done as described by Baleeiro et al.^6^. When applicable, 4.5 kPa ethylene was added to the pressurized culture bottles. For comparison of ethylene and 2-BES, every bottle received 1.7 g L^− 1^ (37 mM) of ethanol in the beginning of the fermentation. 50 mM of 2-BES (sodium salt) was used in one set of duplicates, whereas all bottles without 2-BES received additionally 50 mM NaCl to achieve a similar salinity level. The culture bottles were incubated in a rotary shaker at 37°C and 200 rpm. The pH value was adjusted manually to 5.5 with 4 M KOH or 4 M HCl.

The headspace of the bottles was sampled once or twice per week (depending on methanogenic activity) for monitoring of pressure, gas composition, and gas production. Culture bottles were refilled when their pressure was 130 kPa or lower. The liquid phase was sampled weekly for analysis of organic acids and alcohols. Cell pellets were collected at the end of each batch for microbial community analysis.

### GAS RECIRCULATION REACTORS

Two identical gas recirculation reactor systems were assembled for this study. Each system (Figure 1) consisted of a bioreactor Biostat A plus (Sartorius AG, Göttingen, Germany) with 1.0 L working volume, an 11 L (maximum volume) gas reservoir made of gas-tight, flexible multilayered aluminum-plastic composite material, a condenser, and Hei-Flow Precision peristaltic pumps (Heidolph Instruments GmbH, Schwabach, Germany). The whole system was connected with PVC tubes Tygon^®^ R-3603 or LMT-55 and checked regularly for gas leakages with a gas leak detector GLD-100 (Coy Laboratory Products, Grass Lake, USA). A peristaltic pump ensured a continuous gas flow of ca. 20 mL min^−1^ with injection in the liquid phase through a microsparger. The reactor was operated at 32°C with stirring speed of 300 rpm and 7 kPa overpressure on average. Temperature and pH were monitored and controlled, and oxidation reduction potential (ORP) was monitored.

**Figure 1.**
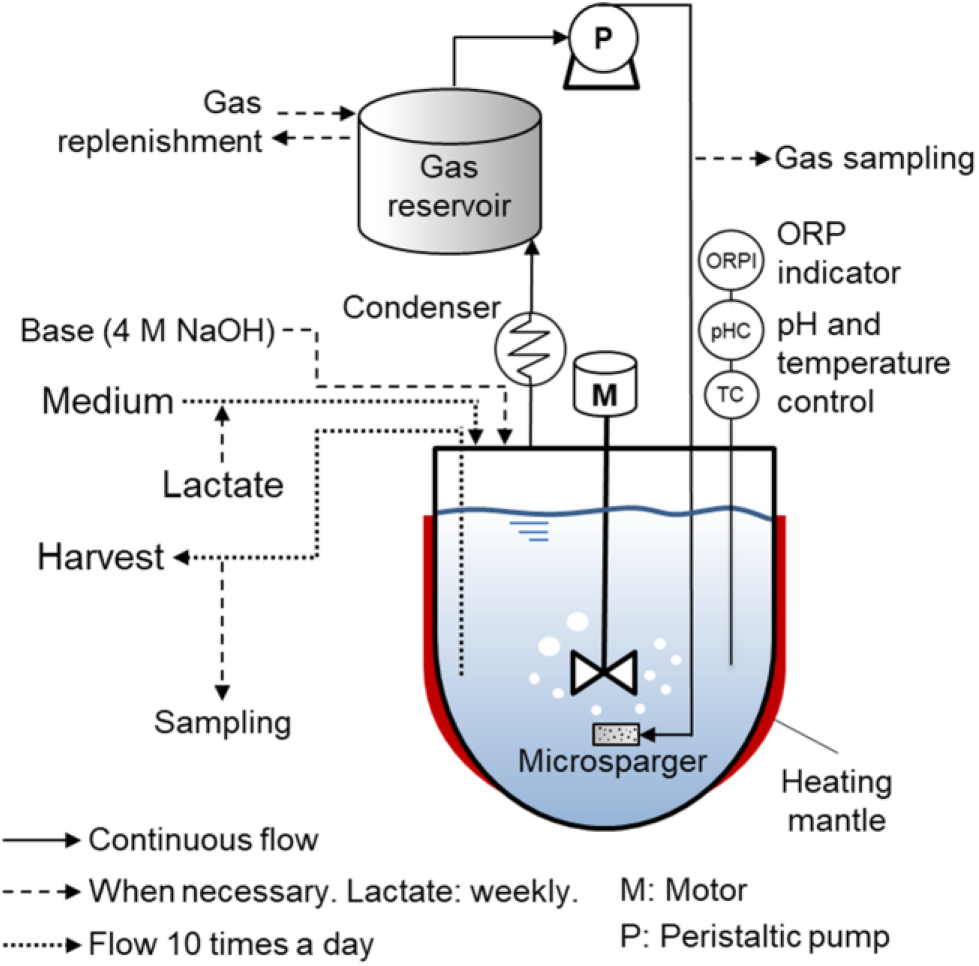
Scheme of the gas recirculation reactor.

The basal medium used in the reactor was similar to the medium used in the culture bottles experiment with the following modifications: it contained 1.61 g L^−1^ NH_4_Cl, did not contain yeast extract, and was prepared with glacial acetic acid instead of a sodium acetate/acetic acid mixture. The basal medium was made anoxic and was sterilized and then stored at room temperature at pH 4.5. The inoculum for the reactor experiment, derived from a biogas reactor, was the same as the one used for “community C” described by Baleeiro et al.^6^, and it was stored under anoxic conditions at room temperature in the dark before its use. For startup, each reactor received 14 vol% of inoculum plus 86 vol% of basal medium with pH adjusted to 6.0. Anoxic, concentrated cysteine and vitamin solution were added to the basal medium immediately before its addition to the vessel or its connection to the feed pump. 4 M NaOH was used to maintain the pH value between 5.9 and 6.0 and as sodium source. Depending on the amount of NaOH added, salinity of the broth was estimated to be in the range of 16 to 28 g L^−1^ NaCl equivalents. Feeding and harvesting were done with peristaltic pumps programmed to operate 10 times per day in pulses lasting 90 s each. Flows were set to a hydraulic retention time (HRT) of 14 days. On day 9 of operation, DL-lactic acid (90% purity) started to be fed with a syringe once a week in order to reach a lactate concentration of 6.0 g L^−1^ (67 mM) after each injection.

Before startup, the gas reservoirs of the assembled dry reactor systems were vacuumed and filled several times with 10 L N_2_ to remove all O_2_. A defined amount of helium was injected in the system and recirculated to estimate the rigid volume of the system (volume of the system without the gas reservoir). After inoculation and for every gas purging/replenishment cycle, the gas reservoir was emptied with a vacuum pump Laboport^®^ N810FT (KNF Neuberger GmbH, Freiburg, Germany) and refilled with 10 L of H_2_:CO_2_ (80:20). Furthermore, 120 mL helium was injected with a syringe to act as an inert tracer gas to quantify volume variations due to microbial activity. Concentration of N_2_ was monitored to identify and quantify possible air contamination in the system. For inhibition of methanogenesis, ethylene was added after gas replenishment to one of the reactors ensuring a minimum ethylene share of 1.3% in the recirculating gas. Once methanogenic activity was established, gas reservoirs were refilled once a week or when the H_2_ share was below 60%, whichever came first. Gas purging and replenishment cycles were always preceded and succeeded by gas sampling in order to keep track of the volume of the system.

The reactor broth was sampled three times per week and before and after each lactate addition. The sampled broth was used for collection of cell pellets for microbial community analysis and for biomass concentration and chemical composition analysis.

Assumptions and calculation steps for the gas recirculation experiment as well as conversion factors used for the electron and carbon balances (Table S2) are described in the Supporting Information.

### ANALYTICAL METHODS

Biomass concentration was determined by measuring the optical density at 600 nm (spectrophotometer Genesys 10 S, Thermo Scientific Inc., Waltham, USA). Concentration of linear monocarboxylates C1-C8, normal alcohols C2-C6, *i*-butyrate, *i-*valerate, *i*-caproate, and lactate was measured by high performance liquid chromatography with a refractive index detector (HPLC Prominence-i RID, Shimadzu Europa GmbH, Duisburg, Germany). H_2_, CO_2_, CH_4_, He, N_2_, and ethylene in the gas phase were analyzed by gas chromatography with a thermal conductivity detector (Light Gas Analyzer ARNL4159 model 4016, PerkinElmer Inc., Shelton, USA). Details of the chromatographic techniques were described previously^6, 18^.

### MICROBIAL COMMUNITY ANALYSIS

Cell pellets collected from serum bottles and from the gas recirculation reactors were washed with phosphate-buffered saline (PBS, 12 mM PO_4_^−3^) solution and stored at −20℃ until their use for amplicon sequencing of the 16S rRNA genes. Detailed description of DNA extraction, PCR, Illumina library preparation, and sequencing on the MiSeq platform can be found in the study of Logroño, et al.^19^. The used primers targeted the V3 and V4 regions of the 16S rRNA gene and were described by Klindworth, et al.^20^. The bioinformatics workflow used for sample inference from amplicon data was described previously6. Taxonomic assignment of amplicon sequence variants (ASVs) was done using the SILVA 138 reference database^21–22^. For the most abundant ASVs, MegaBLAST^23^ was used to find the most similar sequences of cultured species within the NCBI 16S ribosomal RNA sequences database^24^. Phyloseq package for R^25^ was used for filtering, agglomeration, normalization, subsetting, and visualization of the microbiome census data. All samples were rarefied to an equal sequencing depth of 44,017 counts. Raw sequence data for this study was deposited at the European Nucleotide Archive (ENA) under the study accession number PRJEB41050 (http://www.ebi.ac.uk/ena/data/view/PRJEB41050).

### ASSUMPTIONS FOR ECONOMIC ANALYSIS

Assumptions adopted for the economic analysis are described in detail in the Supporting Information.

## RESULTS AND DISCUSSION

The study was divided in two main experiments, namely batch tests in culture bottles and the operation of gas recirculation semi*-*continuous reactors. First, two batch tests were performed in serum bottles to understand the effectiveness of ethylene as an inhibitor, its stability, its effect on the carboxylate production, and to compare it with 2-BES. Afterwards, the gas recirculation reactor system was assembled and operated for 84 days.

To consider chemicals in the gaseous and aqueous phases simultaneously, results are presented in terms of electron equivalents. When relevant, reference is made to results in terms of chemicals concentrations in the Supporting Information.

### INHIBITION OF METHANOGENESIS IN BATCH CULTURES

The electron balances for the first test are shown in Figure S1. Regardless of ethylene presence, more acetate was consumed when H_2_ was present. The presence of H_2_ with ethylene allowed a 3.7-fold higher butyrate and a 4.1-fold higher *i*-butyrate production, as well as a 56% higher caproate production in comparison to cultures with the presence of H_2_ alone (Figure S1-A). 243 ± 2 mmol e^−^ H_2_ was consumed and 249 ± 3 mmol e^−^ CH_4_ was produced when ethylene was not present (Figure S1-B). The presence of ethylene inhibited virtually all methane production (0.1 ± 0.1 mmol e^−^ CH_4_ produced), nevertheless 22 ± 4 mmol e^−^ H_2_ was consumed. Caproate production in this batch experiment was low. Cultures with H_2_ produced slightly more caproate on average (uninhibited: 2.0 ± 2.4 mmol e, with ethylene: 3.2 ± 2.3 mmol e-) than H_2_-free controls (uninhibited: 1.4 ± 1.7 mmol e-, with ethylene: 1.2 ± 0.9 mmol e-) (Figure S1-A). Abiotic controls showed no changes in chemical concentration (data not shown). CH_4_ was not produced in cultures that did not receive exogenous H_2_, indicating that acetoclastic methanogenesis did not play a role.

Afterwards, ethylene and 2-BES were compared in the presence of H_2_, CO_2_, and ethanol in different culture bottles (Figure S2). When H_2_ was present but no inhibitor was used, CH_4_ was the most common product. Both 2-BES and ethylene completely inhibited methanogenesis (Figure S2-B). The use of inhibitors did not strongly affect butyrate or caproate formation (Figure S2-A), however, 2-BES inhibited *i*-butyrate production (0.51 ± 0.07 mmol e^−^).

To account for the possibility of ethylene consumption by the community, the amount of ethylene in the headspace of the bottles was monitored throughout the two batch experiments (Figure S3). No sign of ethylene consumption was found during the periods of the batch experiments. The observed stability of ethylene in the anaerobic cultures is in agreement with a previous study^14^, in which ethylene was not consumed over the whole period of the study (more than 3 months).

A comparison of the microbial community compositions of the inoculum and the inhibited cultures showed that *Methanobacterium* and *Methanobrevibacter* were the methanogens inhibited by ethylene and 2-BES (Figure S4). Being a closed batch system, a complete disappearance of methanogens could not be observed. In the presence of H_2_/CO_2_ and ethylene, the relative abundance of *Clostridium* sensu stricto 12 increased (Figure S4-A). When ethanol was also available, an *Anaerovoracaceae* bacterium had the biggest increase in relative abundance (Figure S4-B). Similar observations were found in a previous study^6^, where *Clostridium* sensu stricto 12 grew most when H_2_ was the only electron donor, but *Anaerovoracaceae* bacteria were most abundant when ethanol was co-fed. The patterns of the community inhibited by 2-BES or ethylene were similar (Figure S4-B). A detailed discussion of the batch experiment results can be found in the Supporting Information.

### COMPONENT BALANCES IN THE GAS RECIRCULATION REACTORS

The operation of the two H_2_/CO_2_ recirculation reactors was divided in two phases lasting 42 days each. The first phase was used for reactor startup and process stabilization. By the end of the start-up phase, hydrogenotrophic methanogenesis was well established in both reactors and ethylene was added to one of the reactors, starting the inhibition phase.

Figure 2 summarizes the results of the two reactor experiments with the profiles of the accumulated compounds during the 84 days of fermentation. Figure S5 presents the concentration profiles of compounds in the aqueous phase. Both reactors were fed with the same amount of lactate, however, the control reactor started lactate consumption later and some lactate was washed out in the beginning (Figure 2).

**Figure 2.**
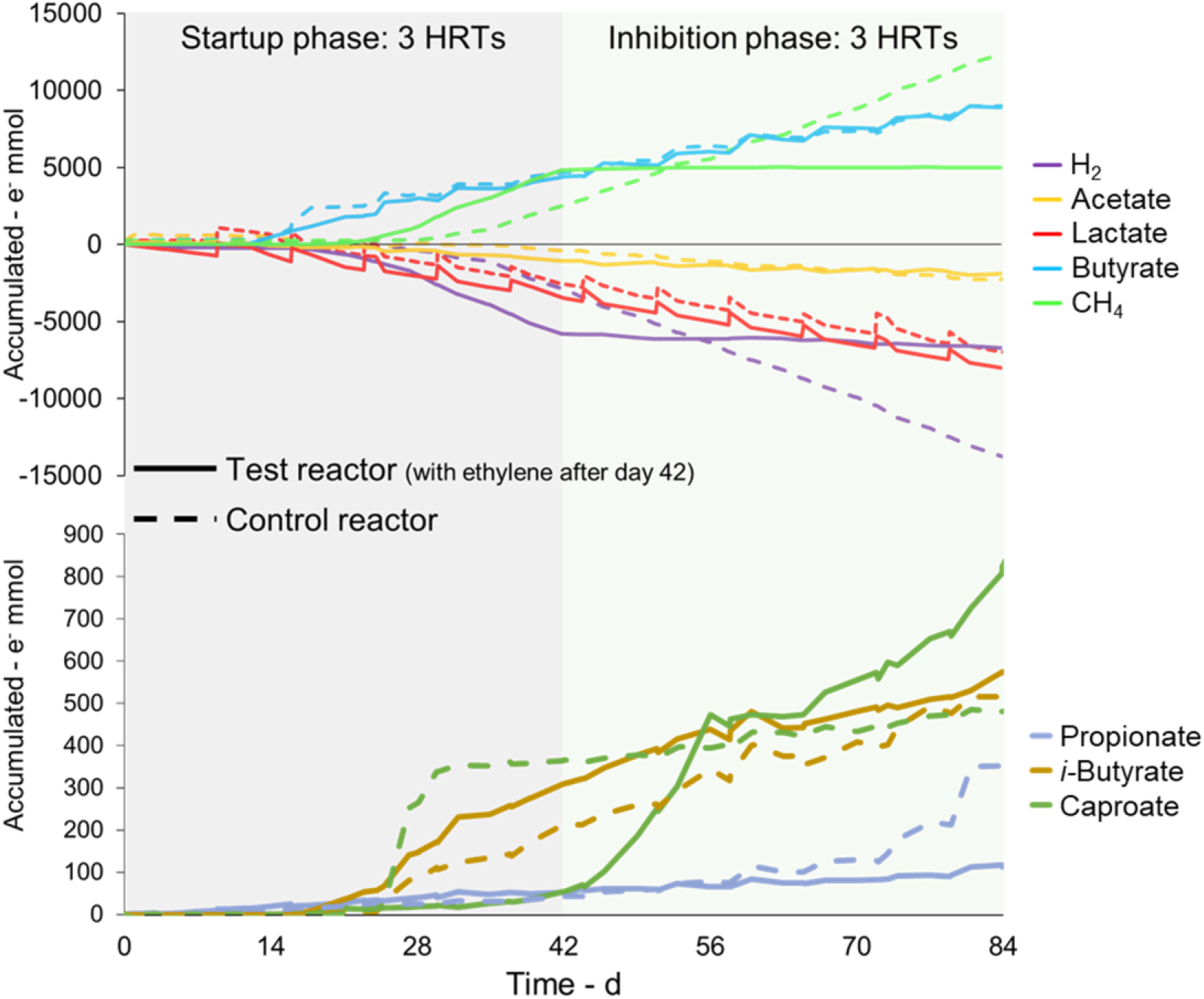
Accumulated substrate consumption and formation of compounds in the reactor with ethylene (Test reactor) and in the reactor without ethylene (Control reactor) shown in electron equivalents. Negative values mean consumption of the compound.

Butyrate was the main carboxylate produced. Weekly spikes of the substrates were reflected by the curves of lactate, butyrate (Figure 2 and Figure S5), and acetate (Figure S5) suggesting that butyrate production was directly coupled with simultaneous consumption of lactate and acetate. Moreover, net consumption of acetate occurred regardless of whether methanogenesis was inhibited or not.

Butyrate production started earlier (day 14) than methanogenesis (between days 21 and 28). CH_4_ formation rates were 216 and 262 mmol e^−^ L^−1^ d^−1^ in the control and in the test reactor, respectively, in the last seven days of the startup phase. CH_4_ production stopped immediately after ethylene addition in the test reactor and H_2_ consumption slowed down from 269 to 23 mmol e^−^ L^−1^ d^−1^. Regarding H_2_ availability, the partial pressure of H_2_ in the control reactor often reached zero and fluctuated strongly in the range of 0 - 80 kPa (Figure S6). In the test reactor, partial pressures of H_2_ and ethylene were within the range of 68 - 80 kPa and 1.3 - 4.8 kPa, respectively. Caproate and propionate production was not clearly related to lactate consumption (Figures 2 and S5). An onset of propionate production, peaking at about 1 g L^−1^, was observed in a late stage in the control reactor despite unchanged operating conditions. *i*-Butyrate production was not inhibited by the use of ethylene and its accumulation remained steady in both the control and the test reactor (Figure 2). In the control reactor, variations of *i-*butyrate concentration followed variations of butyrate concentration (Figure S5-A) while this relation was less clear in the test reactor (Figure S5-B). A discussion on the possible role of metabolic intermediates can be found in the Supporting Information.

The reactor with ethylene showed 55.7% less net consumption of acetate and higher accumulation of electrons in the butyrate and caproate pools (Figure S7-A). Electron selectivity in the test reactor was 101% lactate-to-butyrate (88.2% in the control reactor), 16.9% lactate-to-caproate (2.3% in the control reactor), and 1.4% lactate-to-propionate (6.3% in the control reactor). Channeling of H_2_ to CH_4_ was mainly responsible for the consumption of 10.9 moles e^−^ H_2_ in the control reactor and less than 10% of this consumption (0.93 mol e^−^ H_2_) was observed in the reactor that received ethylene (Figure S7-B). Despite being a small amount of electrons in comparison to the methanogenic process, the H_2_ consumption in the reactor with ethylene accounted for 17% of the total electron donor consumption. In contrast, the non-methanogenic H_2_ consumption accounted for 6.6% of the total electron donors consumed in the control reactor.

### NET CARBON FIXATION

Both reactors started with net carbon fixation, but there was a trend in the long run for loss of carbon fixation capacity (Figure S8). The test reactor became a net carbon emitter by the 3^rd^ HRT period, whereas the control reactor became a net carbon emitter in the 5^th^ HRT. The use of ethylene, injected for the first time in the beginning of the 4^th^ HRT, reverted the trend for the test reactor (Figure S8). A maximum carbon fixation rate of 62.2 mmol C per 14-days period was observed, which was equivalent to 0.20 g CO_2_ L^−1^ d^−1^. CO_2_ dissolution in the broth had only a small impact in the carbon fixation estimation.

### MICROBIAL COMMUNITY DEVELOPMENT

The development of the microbial communities in the gas recirculation reactors is shown in Figure 3. By the end of the startup phase (day 42), the acidogenic genera *Clostridium, Caproiciproducens, Eubacterium,* and *Oscillibacter*, together with the methanogenic genus *Methanobacterium*, were the main settlers in both reactors. *Bacteroides* settled in both reactors from the middle until the end phase of the experiment. *Rummeliibacillus*, *Sutterella*, *Defluviitoga*, *Fastidiosipila*, and unclassified *Actinomycetaceae* were only transiently detected during the startup phase.

**Figure 3.**
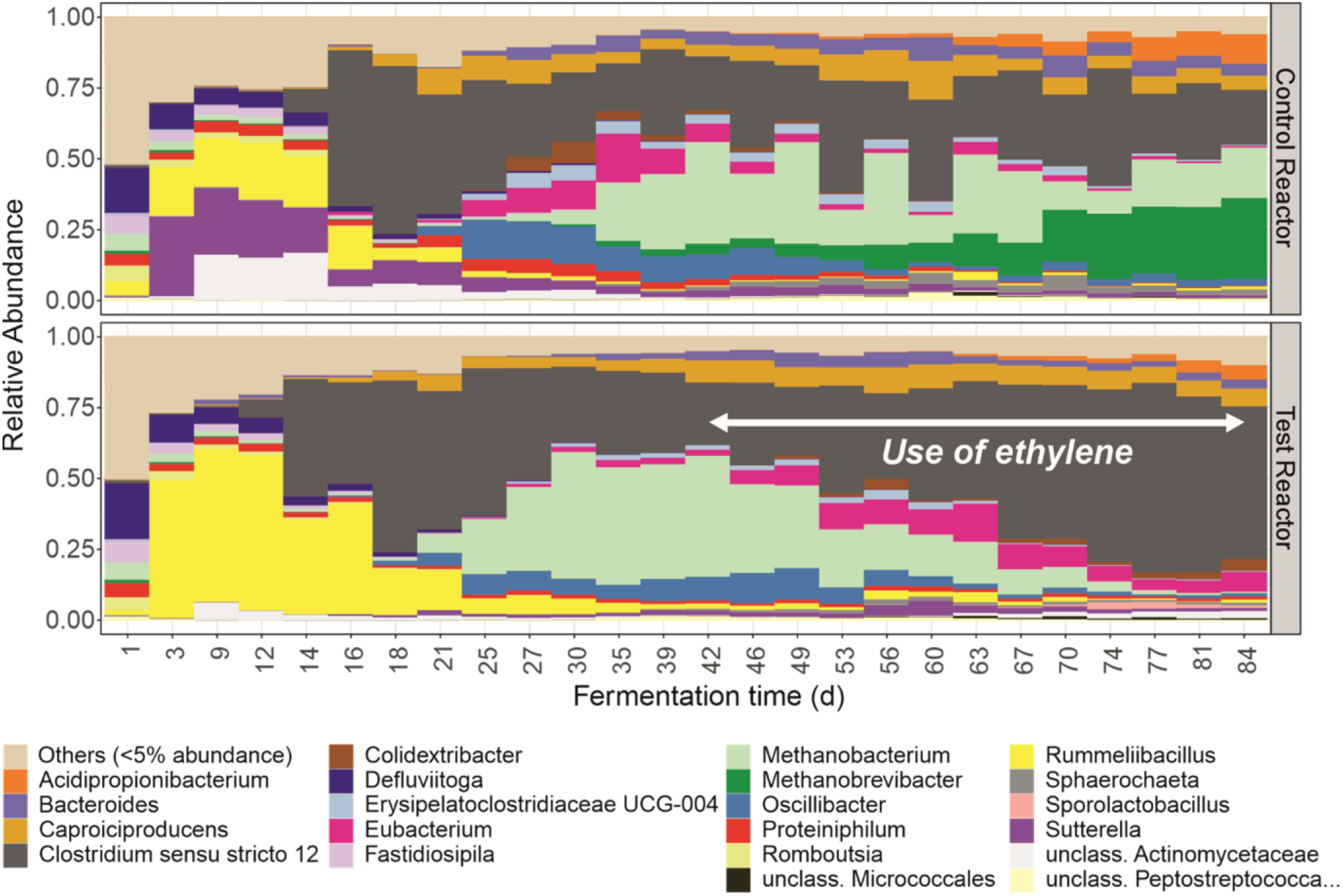
Microbial community profiles for the control and the test reactors. The latter received ethylene after day 42. Hydrogenotrophic methanogens of the genus *Methanobacterium* were washed out during the period in which ethylene was used.

In the control reactor, an additional methanogenic genus, *Methanobrevibacter*, ascended in the later experimental phase while *Eubacterium* and other less abundant genera were washed out from the reactor. With the use of ethylene in the test reactor, the washout of *Methanobacterium* gave room for higher relative abundances of *Clostridium*, *Eubacterium*, and *Colidextribacter*, genera that harbor acetogenic and acidogenic species. A slow but steady increase of *Acidipropionibacterium* was observed in the late fermentation stages in particular in the control reactor (Figure 3), which temporally corresponds to the profile of propionate concentration in the same period (Figure S5-A).

Butyrate formation was positively correlated with the relative abundances of *Clostridium* sensu stricto 12 while caproate correlated positively with abundances of *Eubacterium*, *Oscillibacter*, *Caproiciproducens*, *Erysipelatoclostridiaceae* UCG-004, and *Colidextribacter* (p<0.01, Figure S9). Production of CH_4_ correlated positively with relative abundances of *Methanobacterium* and *Methanobrevibacter* (Figure S9), known as hydrogenotrophic methanogens. *i-*Butyrate production correlated with abundances of *Oscillibacter*, *Caproiciproducens*, *Bacteroides*, and *Erysipelatoclostridiaceae* UCG-004 (Figure S9). Presence of ethylene correlated with higher relative abundances of *Clostridium* sensu stricto 12, *Eubacterium*, *Caproiciproducens*, *Colidextribacter*. It is worth notice that the analysis shows no distinctions between direct and indirect correlations. This is exemplified in the cases that are clearly indirect: correlations between abundances of some bacterial genera and CH_4_ formation and between methanogens and formation of propionate and *i-*butyrate. Some of the ASVs within the acidogenic and acetogenic genera which had high similarities with isolates are presented in the Supporting Information.

Although the planktonic methanogens were observed to be almost completely washed out with the use of ethylene in the test reactor, biofilms attached to the vessel walls and other inner reactor surfaces still contained methanogens at the end of the experiment (Figure S10).

### ETHYLENE AS A SCALABLE INHIBITOR

While 2-BES can be considered a specialty chemical (41 US$/kg), ethylene is a commodity with a relatively low price (1 US$/kg) and widely available on the chemical market. Moreover, ethylene (as well as acetylene) is a common constituent of syngas from biomass gasification in the concentrations used in this study^26–27^. With gas recirculation, ethylene could be used as a recyclable methanogenesis inhibitor, which is not the case for 2-BES, as the latter is soluble in water and would be washed out from the aqueous phase. On the other hand, recirculation of gas increases the auxiliary power requirement of the process. Figure 4 presents an order of magnitude estimate of the operating costs per m^3^ of broth for gas recirculation (depending on the compression pressure) and for the use of 2-BES (depending on its concentration). As a reference for economic feasibility, the potential value that can be obtained by selling the carboxylates present in the fermenter broth is estimated to be between 8 US$ m^−3^ and 40 US$ m^−3^ (Figure 4). This value depends on the extractable carboxylate composition and the selling price of the carboxylates. Assumptions adopted for the economic analysis are described in detail in the Supporting Information. Four alternatives were compared: i) H_2_/CO_2_/ethylene recirculation at the flow used in this study (1.2 L h^−1^ L^−1^); ii) H_2_/CO_2_/ethylene recirculation at a flow at optimized conditions with ten times the microbial gas consumption observed in the test reactor with ethylene (0.14 L h^−1^ L^−1^); iii) recirculation of ethylene (0.018 L h^− 1^ L^−1^); and iv) use of 2-BES for methanogenesis inhibition at concentrations up to 50 mM (10.5 g L^−1^ sodium 2-BES). The gas recirculation alternatives i)-iii) cost between 0.02 US$ m^−3^ and 2 US$ m^−3^, which is well below the value range of the carboxylate broth. The use of 2-BES for inhibiting methanogenesis (option iv)) costs at least 43 US$ m^−3^ and is thus not economically feasible even at concentrations below those required for inhibition of hydrogenotrophic methanogens. For a more detailed economic analysis in the future, the supply of H_2_ and CO_2_ as well as the pay-offs of gas recirculation (in terms of increased selectivity to MCC, higher carboxylate production, and net carbon fixation) would have to be considered. As a reference, an encompassing economic analysis considering downstream processing and capital costs (but excluding gas recirculation and methanogenesis inhibition costs) was done previously by Scarborough, et al.^28^ for an integrated lignocellulosic biorefinery producing MCC, ethanol, and electricity.

**Figure 4.**
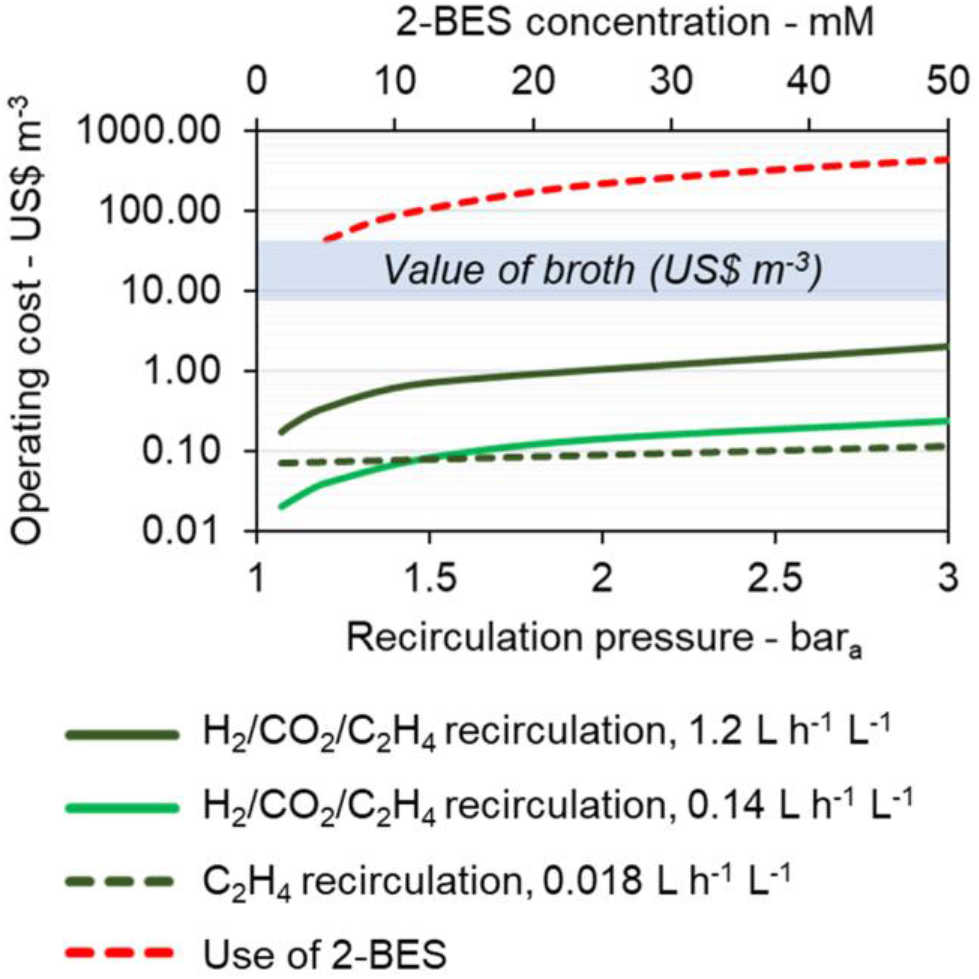
Estimated operating costs of 2-BES addition or H_2_, CO_2_, and ethylene (C_2_H_4_) recirculation depending on the pressure. The value of the carboxylate-containing broth is estimated to be between 8 US$ m^−3^ and 40 US$ m^−3^.

Another difference between ethylene and 2-BES found in the batch experiments was that 2-BES showed inhibitory effects on *i*-butyrate formation, whereas ethylene did not. This fact could prove useful if the production of branched carboxylates is desired, in particular considering that *i*-butyrate and *i*-caproate have been recognized as potential bio-product platforms^29–30^.

As a cautionary tale, it was shown that the biofilm formed in inner reactor parts contained methanogens from previous reactor operation phases (Figure S10). Since ethylene is a reversible inhibitor^14^, the planktonic community may be easily re-inoculated with methanogens from the biofilm or unsterile substrate if the use of ethylene ceases.

### MECHANISM OF ETHYLENE INHIBITION

Research on the mechanism of ethylene inhibition of methanogens and its effects on acidogenic bacteria has been not as encompassing as research with acetylene. Arguably, the mechanism of inhibition by ethylene may be comparable to that of acetylene, since ethylene also has a π C-C bond^31^. The specific inhibition by ethylene may be explained by its effect on some types of hydrogenases, specifically those which methanogens most depend on31. Acetylene was shown to strongly inhibit [NiFe] hydrogenases of a methanogen (*Methanosarcina* sp. MST-AI DSM 2905) and of a sulfate-reducing bacterium (*Desulfovibrio gigas*) while presenting no effect on the nickel-free hydrogenase of another sulfate^−^ reducing bacterium (*Desulfovibrio vulgaris*)^32^. Methanogenic archaea depend on [NiFe] hydrogenases for fast H_2_ oxidation^33^ whereas fermentative bacteria (in particular *Firmicutes*) have a vast diversity of [FeFe] hydrogenases^34^. Whether ethylene has a differential effect on these two types of hydrogenases needs to be tested in future studies, in particular because alternative inhibition mechanisms have been proposed^35–36^. It is worth noticing that the effects of acetylene and ethylene on anaerobic cultures are not identical. Acetylene is a less selective inhibitor than ethylene since acetylene inhibits sulfate-reducing and nitrogen-fixing bacteria whereas ethylene does not^14,37^. Ethylene also seems to be more stable in anaerobic systems than acetylene, since a rare metabolic pathway has been found that degrades acetylene in the absence of strong electron acceptors (e.g. sulfate)^38–39^ while to the best of our knowledge no similar pathway is known for ethylene degradation.

### POSSIBILITIES FOR PROCESS OPTIMIZATION

Developing, controlling, and optimizing a mixotrophic acidogenic community is not trivial because homoacetogens typically prefer higher ATP-yielding substrates (e.g. carbohydrates, ethanol, lactate) before switching to autotrophic metabolism, as seen in the case of *C. ljungdahlii* in the presence of fructose^40^. Under lactate starvation, homoacetogens are forced to put their substrate flexibility into use^41^. Here, the reactor system was operated in such a way that H_2_ and CO_2_ were always available, basal medium (with acetate) was fed ten times per day, and lactate was fed once a week. It is possible that this feeding strategy, which had a longer intermittency for lactate, may have helped favor autotrophy over heterotrophy. Another possible consequence of longer intervals of substrate feeding is the maintenance of high diversity in the community^42^. High community diversity is a factor that can help couple non-methanogenic H_2_ consumption with MCC formation^6^.

Acetate and lactate were used as a simplified model of an ensiled feedstock or the organic fraction of municipal solid waste. The exploration of the concept with complex biomass is the next step to start assessing the economic feasibility of the H_2_, CO_2_, and ethylene gas recirculation approach. Future studies should also aim to increase caproate concentration, since in this study the maximum caproate concentration achieved (up to 1.2 g L^−1^, Figure S5) fell short in comparison to state-of-the-art chain elongation processes. Furthermore, increase of non-methanogenic gas consumption rates is highly desirable, since the non-methanogenic H_2_ consumption achieved here was only 17% of the total electron donor consumption. In this direction, operation with CO or syngas mixtures may help improve gas consumption, chain elongation, solventogenesis, and volumetric rates^43^. Besides, the presence of CO can help inhibit methanogens^44^.

Depending on the desired products (SCC, MCC, alcohols, *i*-butyrate, etc.), the operation of the system may be optimized by adjusting gas-liquid feeding strategies, together with other operational parameters such as pH and temperature. Nevertheless, a better knowledge of the relation between process parameters and production of SCC, MCC, and alcohols is still needed.

## ASSOCIATED CONTENT

### Supporting Information

The following files are available free of charge.

Table S1 and S2, Figures S1 to S10, calculations for component balances, assumptions for economic analysis, further discussion of the results from the batch experiments, possible metabolic intermediates, and ASV similarities to species level. (PDF)

## AUTHOR INFORMATION

### Author Contributions

FCFB, SK, and HS conceptualized the study and reviewed the manuscript. FCFB developed the methodology, performed the experiments, analyzed the data, and prepared the original draft. HS and SK supervised the project and supported the data analysis. All authors have given approval to the final version of the manuscript.

## ACKNOWLEDGEMENT

We thank Ute Lohse for technical assistance in library preparation for MiSeq amplicon sequencing as well as Carolin Köbe and Felix Kaufmann for helping with the batch experiments. The DBFZ Department Biorefineries is acknowledged for providing the reactors.

## SYNOPSIS

Anaerobic fermentation with continuous recirculation of H_2_, CO_2_, and ethylene increases selectivity to carboxylates while allowing net carbon fixation.

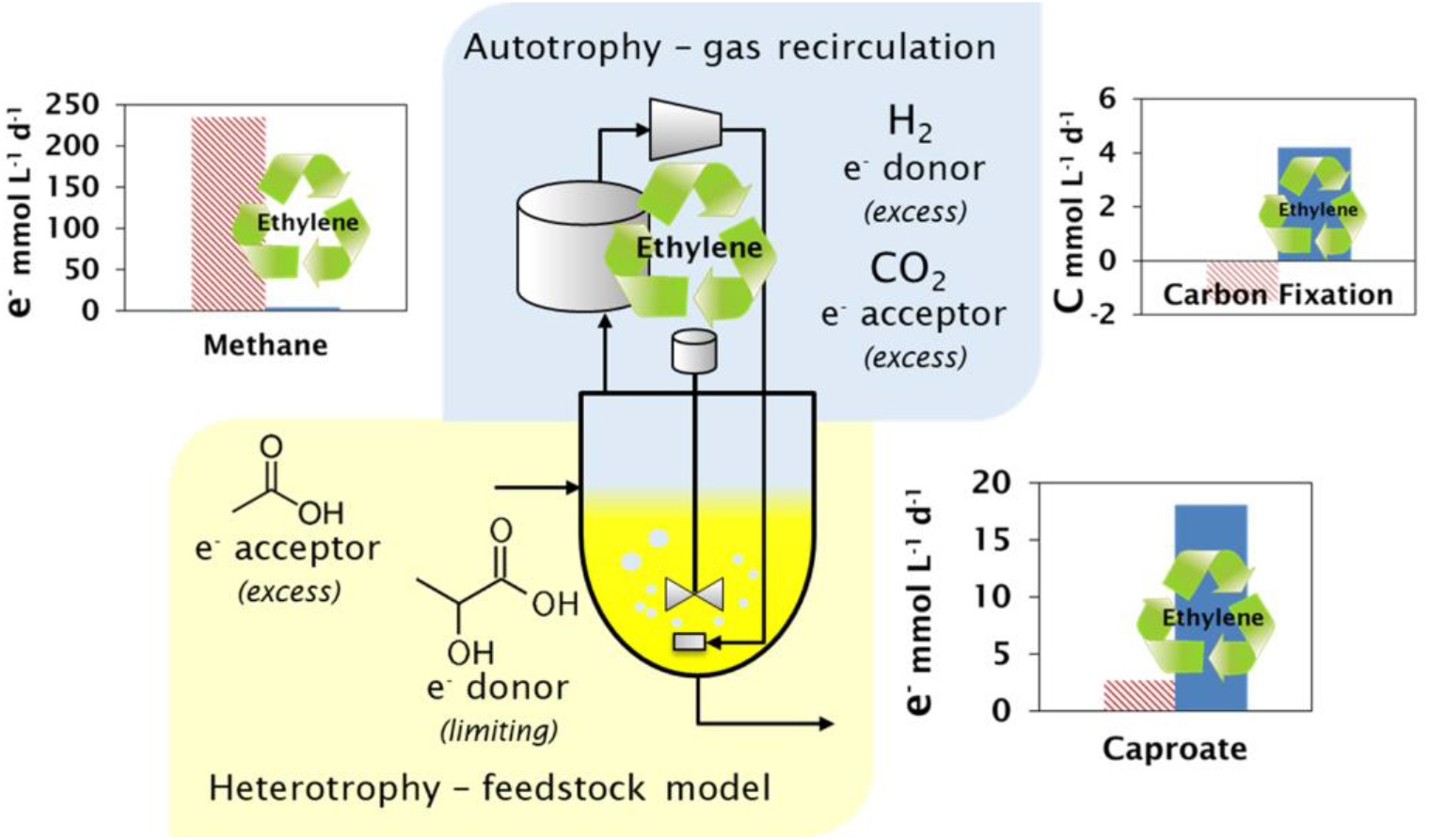
For Table of Contents Only.

