## Supporting Information for "Recirculation of H_2_, CO_2_, and ethylene improves carbon fixation and carboxylate yields in anaerobic fermentation"

Table S1. 2
Table S2. 2
Calculations for component balances. 3
Assumptions for economic analysis 4
Figure S1. 5
Figure S2. 6
Figure S3. 6
Figure S4. 7
Further discussion of the results from batch experiments. 7
Figure S5. 9
Possible metabolic intermediates. 10
Figure S6. 10
Figure S7. 11
Figure S8. 11
Figure S9. 12
Figure S10. 13
ASV similarities to species level. 14
References 14

**Table S1.** Fermentation conditions tested in the batch experiments.

| *Experiment* | $H_2$:$CO_2$ (Effect of ethylene and stability) | | | $H_2$:$CO_2$ + 1.7 g $L^{-1}$ ethanol (Ethylene vs. 2-BES) | | |
|---|---|---|---|---|---|---|
| *Condition* / *Inhibitor* | **$H_2$:$CO_2$** | **$N_2$:$CO_2$** | **Abiotic** | **$H_2$:$CO_2$** | **$N_2$:$CO_2$** | **Abiotic** |
| Uninhibited | 2 bottles | 2 bottles | 1 bottle | 2 bottles | 2 bottles | - |
| With ethylene | 2 bottles | 2 bottles | 1 bottle | 2 bottles | - | - |
| With 2-BES | - | - | - | 2 bottles | - | - |

**Table S2.** Conversion factors for electron balances.

| Compound (formula) | Molar mass (g $mol^{-1}$) | mol C/mol | mol $e^-$/mol |
|---|---|---|---|
| Biomass ($C_1H_{1.8}O_{0.5}N_{0.2}$) | 24.6 | 1.0 | 4.2 |
| Formic acid/formate ($C_1H_2O_2$) | 46.0 | 1.0 | 2.0 |
| Acetic acid/acetate ($C_2H_4O_2$) | 60.0 | 2.0 | 8.0 |
| Ethanol ($C_2H_6O$) | 46.0 | 2.0 | 12.0 |
| Propionic acid/propionate ($C_3H_6O_2$) | 74.0 | 3.0 | 14.0 |
| Lactic acid/lactate ($C_3H_6O_3$) | 90.0 | 3.0 | 12.0 |
| Butyric acid/butyrate ($C_4H_8O_2$) | 88.0 | 4.0 | 20.0 |
| *i*-Butyric acid/*i*-butyrate ($C_4H_8O_2$) | 88.0 | 4.0 | 20.0 |
| Butanol ($C_4H_{10}O$) | 74.0 | 4.0 | 24.0 |
| Valeric acid/valerate ($C_5H_{10}O_2$) | 102.1 | 5.0 | 26.0 |
| Caproic acid/caproate ($C_6H_{12}O_2$) | 116.1 | 6.0 | 32.0 |
| Caprylic acid/caprylate ($C_8H_{16}O_2$) | 143.1 | 8.0 | 44.0 |
| $H_2$ | 2.0 | 0.0 | 2.0 |
| $CO_2$ | 44.0 | 1.0 | 0.0 |
| $CH_4$ | 16.0 | 1.0 | 8.0 |

CALCULATIONS FOR COMPONENT BALANCES. To enable component balances in the gas recirculation system, a set of calculation steps was done. For the gaseous phase during a sampling procedure, *N*:

$$V_{Total_N} = \frac{y_{He_{N-1}}}{y_{He_N}} V_{Total_{N-1}} \quad (1)$$

$$V_{Gas\ Bag_N} = V_{Gas\ Bag_{N-1}} + \left(V_{Total_N} - V_{Total_{N-1}}\right) \quad (2)$$

where $V_{Total}$ is the total gaseous volume of the system; $y_{He}$ is the molar or volume fraction of the tracer gas, helium; and $V_{Gas\ Bag}$ is the volume of gas in the gas reservoir.

The accumulated gas production for a gaseous compound *i*, $M_i$, on a molar or mass basis was calculated as follows:

$$M_{i_N} = M_{i_{N-1}} + \left(m_{i_N} - m_{i_{N-1}}\right) \quad (3)$$

where $m_i$ is the total amount of compound *i* in the system on a molar or mass basis.

If the sampling is just after the gas replenishment the volume of the gaseous phase is calculated as follows:

$$V_{Gas\ Bag_N} = V_{H2} + V_{CO2} + V_{He} + V_{C2H4} \quad (4)$$

$$V_{Total_N} = V_{Gas\ Bag_N} + V_{rigid} \quad (5)$$

where $V_{H2}$, $V_{CO2}$, $V_{He}$, $V_{C2H4}$ are the volumes of newly injected $H_2$, $CO_2$, He, and $C_2H_4$, respectively; and $V_{rigid}$ is the volume of the gas phase of the system outside the gas reservoir.

There is a discontinuity in the amount and composition of the gas in the system after gas replenishment. To account for it, the gas phase was sampled just before and immediately after the replenished gas mixture is homogenized. Since the gas replenishment and mixing lasts only a short period of time in comparison to the whole duration of the fermentation, it was assumed that

$$t_N \approx t_{N-1} \quad \therefore \quad M_{i_N} \approx M_{i_{N-1}} \quad (6)$$

where $t_N$, in days, is the fermentation time related to the sampling point *N*, just after gas replenishment and homogenization.

For the accumulated balances of compounds in the aqueous phase, both feeding and washing out of chemicals were considered. The washout of chemicals in the broth was estimated using the average of the concentration values at sampling points *N* and *N-1*. Accumulated amounts for the compound *j* at sampling *N* was calculated as follows:

$$M_{j_N} = M_{j_{N-1}} + \left(\frac{c_{j_N} - c_{j_{N-1}}}{2} - c_{j_{feed}}\right) V_{broth} \, \frac{(t_N - t_{N-1})}{HRT} \quad (7)$$

where $M_j$ is the accumulated amount of the aqueous-phase compound *j* on a molar or mass basis; $c_j$ and $c_{j\,feed}$ are the concentrations of compound *j* in the broth and in the feed medium, respectively; $V_{broth}$ is the working volume of the reactor; and *HRT* is the hydraulic retention time, in days.

The amount of carbon fixed by the system during a certain period, $n_{fixed}$, in mmol C, was calculated as follows:

$$n_{fixed} = -(\Delta n_{CH4} + \Delta n_{CO2}) \quad (8)$$

where $\Delta n_{CH4}$ and $\Delta n_{CO2}$ are $CH_4$ and $CO_2$ produced in the period, respectively, in mmol C.

ASSUMPTIONS FOR ECONOMIC ANALYSIS. To estimate operating costs of using 2-BES, a price of 41 US$ per kilogram of sodium 2-BES (cost insurance & freight (CIF) Hamburg) was assumed. The price is based on a quote from a chemicals provider (Changzhou, China) for a purchase of at least 1,000 kg sodium 2-BES with ≥98% purity (CAS Registry Number: 4263-52-9). To estimate costs for acquiring ethylene, an approximate price of 1 US$ $kg^{-1}$ was considered[1] and a continuous loss of 1% of the flow of ethylene was assumed. For electricity consumption by gas recirculation, an efficiency of 90% of an isentropic compressor with an inlet gas pressure of 1.0 $bar_a$ and an electricity price of 0.12 US$ $kWh^{-1}$ was assumed[2]. For the mixture of $H_2$/$CO_2$/ethylene, a gas mixture with 78.5% $H_2$, 20% $CO_2$, and 1.5% ethylene was considered. An HRT of 14 days was considered. For the range of values of the carboxylate broth, an average price of carboxylates of 2 US$ $kg^{-1}$ and an extractable amount of carboxylates between 4 g $L^{-1}$ (lowest broth value) and 20 g $L^{-1}$ (highest broth value)[2] was assumed. Other operating costs of the process, such as power input for stirring, consumption of other chemicals, and downstream processing, were not considered.

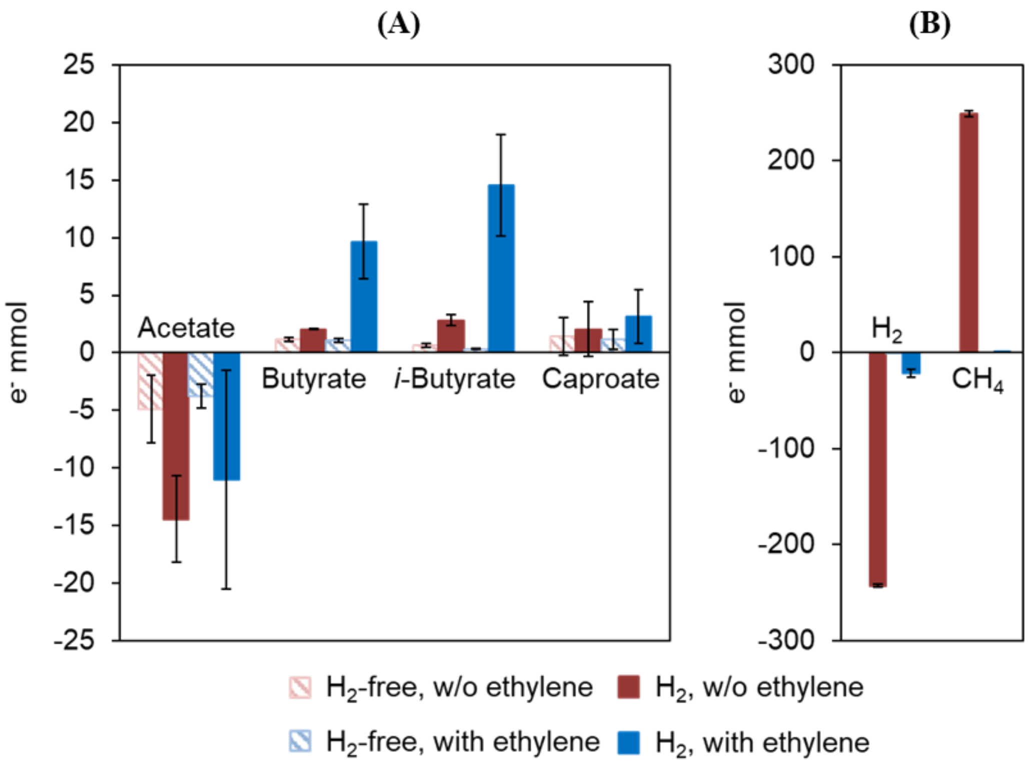


**Figure S1.** Effect of ethylene as inhibitor in an enrichment culture growing on $H_2/CO_2$ in the first batch test. Consumption and production of chemicals in the aqueous phase (A) and in the gas phase (B) after 48 days of fermentation in terms of electron equivalents is shown. $H_2$-free cultures and cultures without ethylene served as controls. Error bars are standard errors.

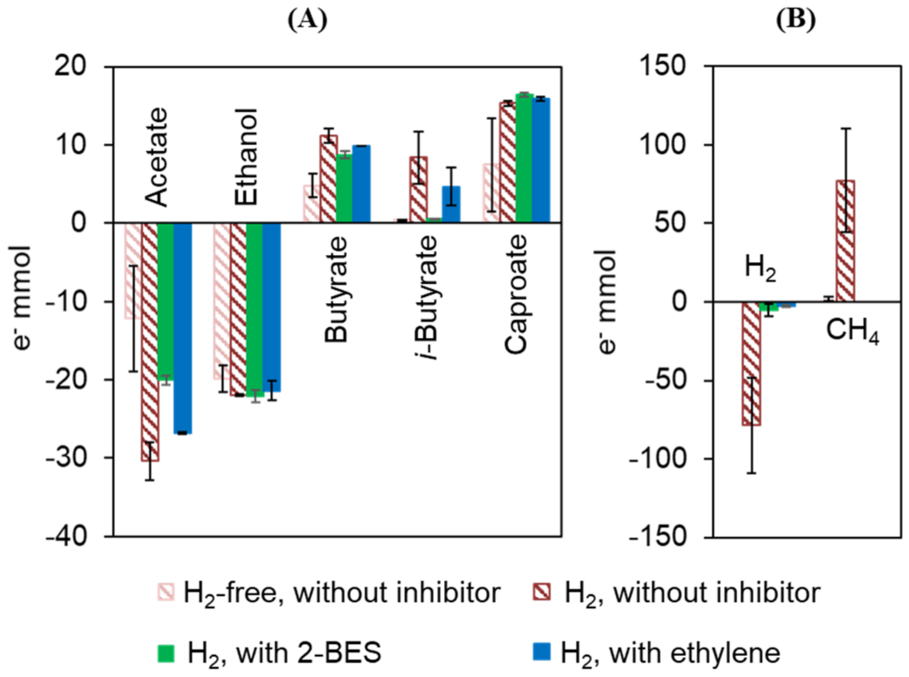


**Figure S2.** Effects of ethylene and 2-BES on cultures fed with $H_2$, $CO_2$, and ethanol in the second batch test. Consumption and production of chemicals in the liquid phase (A) and in the gas phase (B) after 63 days of fermentation in terms of electron equivalents. Error bars are standard errors.

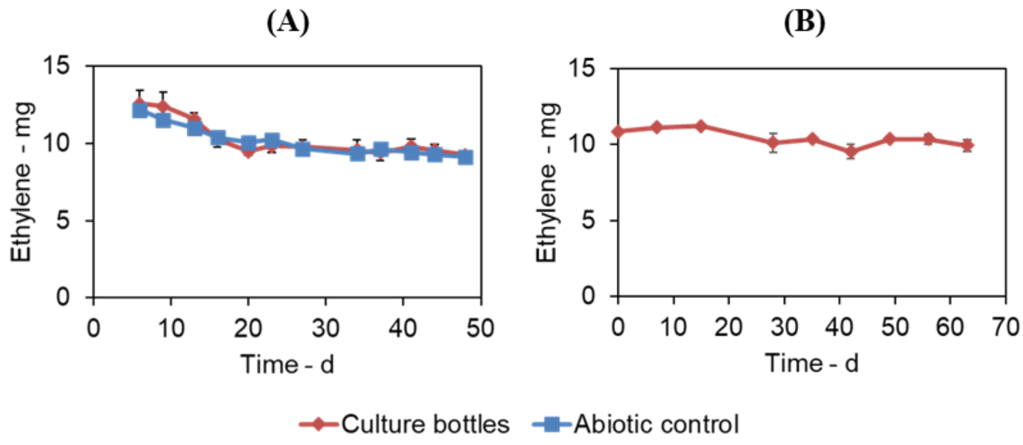


**Figure S3.** Monitoring of the amount of ethylene in the headspace of the culture bottles in the batch experiments. No signs of ethylene degradation were detected. The monitoring was done in the presence of $H_2$ and $CO_2$ with abiotic controls (A) and in the presence of $H_2$, $CO_2$, and ethanol without abiotic controls (B).

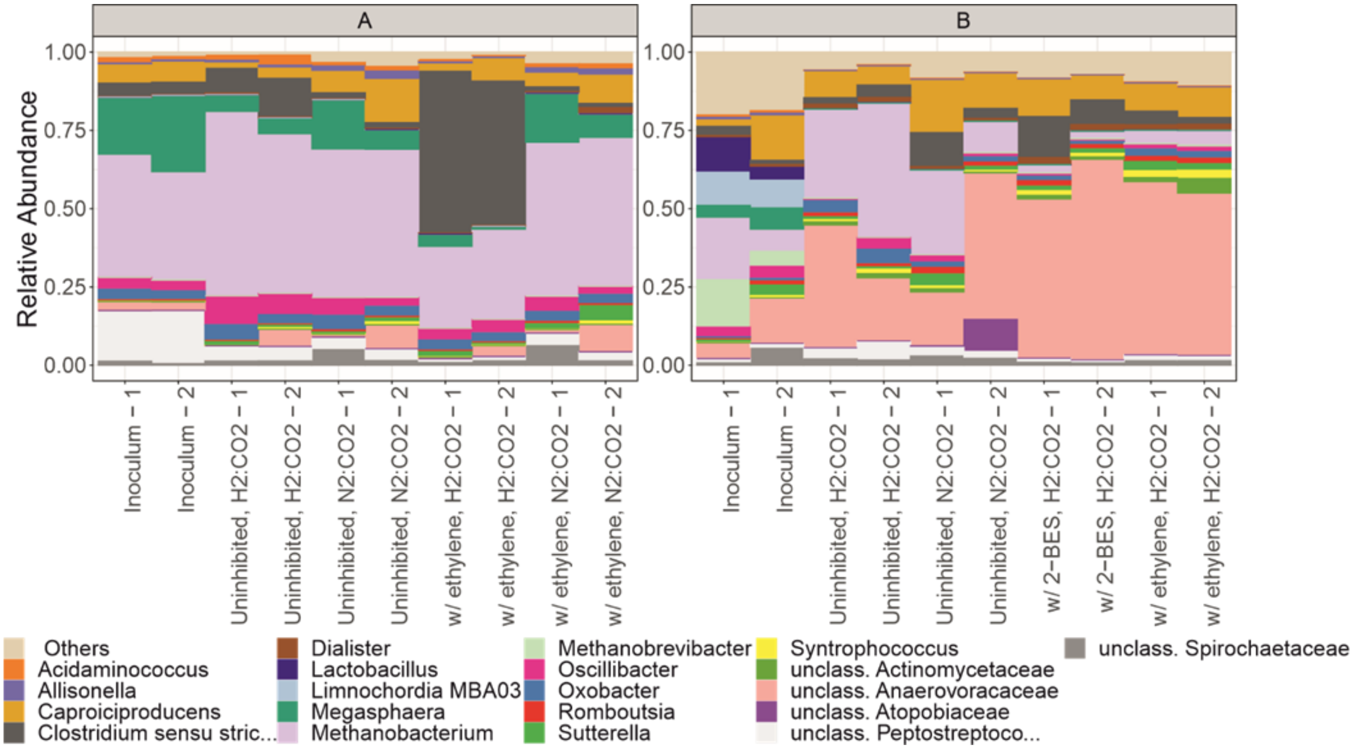


**Figure S4.** Effect of ethylene inhibition on the microbial community composition in the first (A) and second (B) batch experiment. Duplicates are shown and indicated as -1 and -2. The 20 most abundant genera are shown. Abiotic cultures did not show growth and were not analyzed.

FURTHER DISCUSSION OF THE RESULTS FROM BATCH EXPERIMENTS. Communities in the two batch experiments (Figure S4) differed due to slightly different inocula origins. The first batch was inoculated from cultures of the experiment "use of inhibitors" and the second batch was inoculated from the cultures of the experiment "acetate concentration" described by Baleeiro et al.[3]. Both cultures belonged to "community C" and the community of the "use of inhibitors" experiment represented a further enrichment of the community of the "acetate concentration" experiment.

In the first batch experiment of the current study, $H_2$ was the only electron donor given as substrate and little growth was observed when methanogenesis was inhibited (OD600 0.35 ± 0.02 in comparison to 1.2 ± 0.1 when no inhibitor was used). Combined with the fact that the culture bottle was a closed system, the low growth explains why relative abundance of methanogens was still relatively low by the end of the batch in the presence of $H_2$:$CO_2$ and ethylene (Figure S4-A). When ethanol was co-fed with $H_2$ in the second batch (Figure S4-B), stronger growth was observed when methanogenesis inhibitors were present (OD600 0.59 ± 0.03 with ethylene and 0.59 ± 0.02 with 2-BES) and a lower relative abundance of methanogens was observed by the end of the experiment.

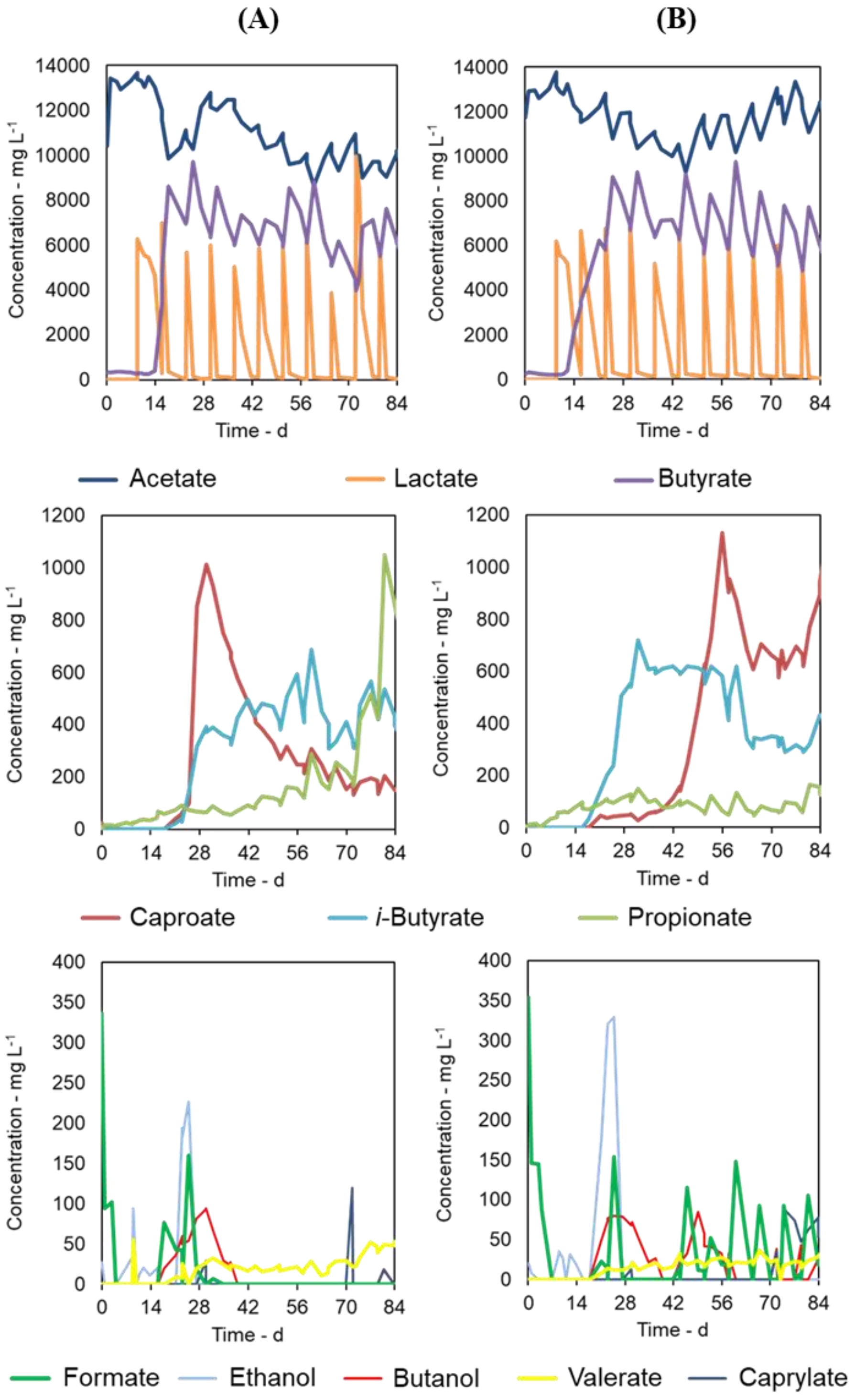


**Figure S5.** Concentration profiles of substrates and metabolites in the control reactor (A) and in the reactor with ethylene addition after day 42 (B).

POSSIBLE METABOLIC INTERMEDIATES. Formate, ethanol, and butanol were detected in low concentrations (<350 mg $L^{-1}$) in appearance/disappearance cycles. Disappearance of these compounds could not be explained solely by washing-out, pointing to the possible role of these compounds as intermediates (Figure S5). After day 42, cycles of butanol and formate continued to occur in the test reactor (Figure S5-B) whereas no more cycles were seen in the control reactor in the same period (Figure S5-A). Formate is an intermediate of homoacetogenesis and can be used as a substrate by some chain-elongating bacteria[4-5]. Short-chain alcohols (methanol, ethanol, and propanol) are electron donors for chain elongation[6]. Butanol, however, has not yet been shown to be an electron donor for chain elongation. Valerate and caprylate concentrations remained below 50 mg $L^{-1}$ and 120 mg $L^{-1}$ during the fermentation, respectively.

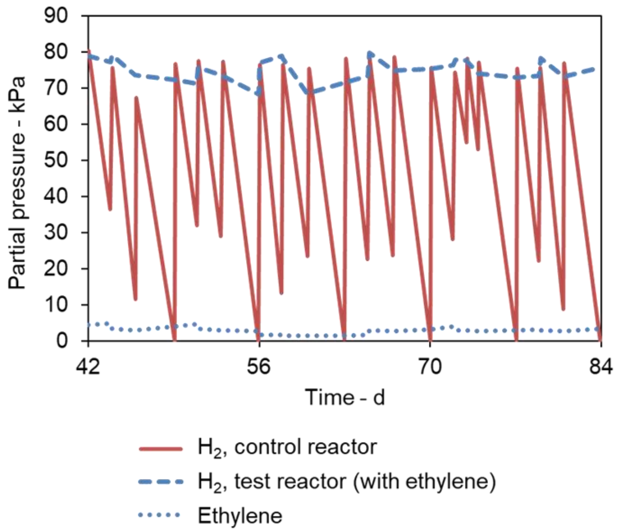


**Figure S6.** Partial pressures of $H_2$ and ethylene in the gas recirculation reactors in the period between operation days 42 and 84. Variations of the partial pressure of ethylene were caused by manual gas purging and refilling procedures. No consumption of ethylene was observed.

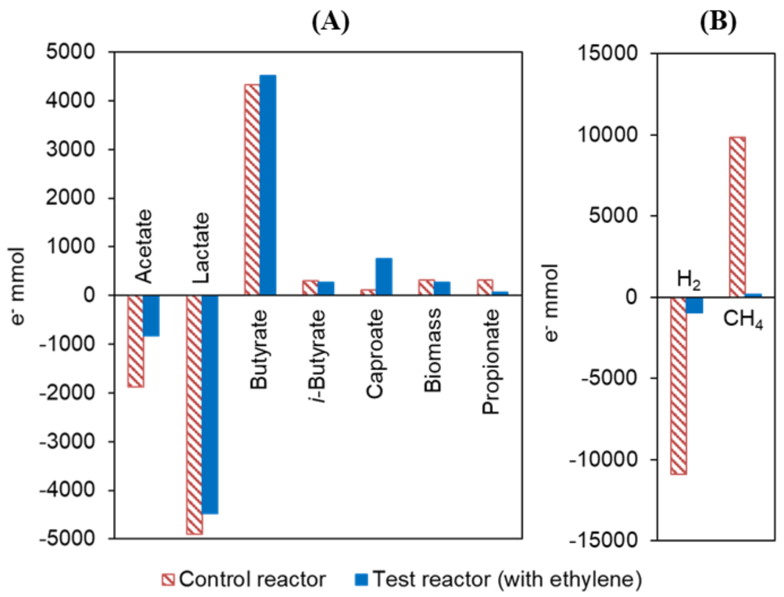


**Figure S7.** Component balances for both reactors in the period when ethylene was used in the test reactor (day 42 to 84) shown as electron equivalents. Production (positive) and consumption (negative) of components in the aqueous phase (A) and in the gas phase (B).

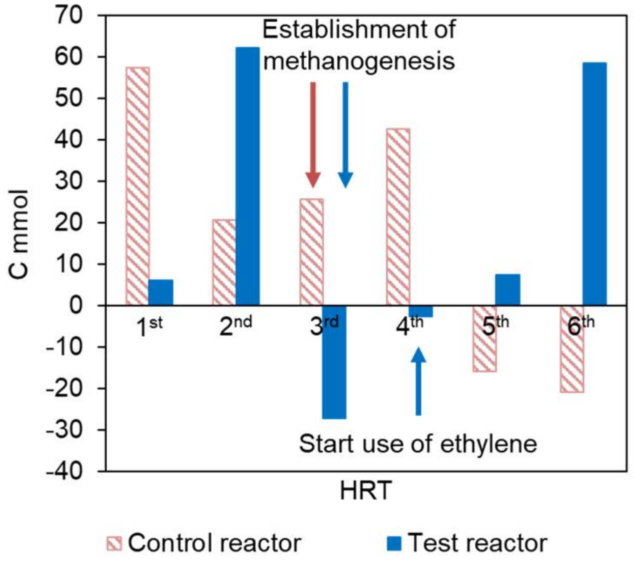


**Figure S8.** Net carbon fixation during each HRT period. One HRT period equals 14 days. The time points of starting methanogenesis in both reactors and of ethylene use in the test reactor are indicated. Assuming a $CO_2$ partial pressure of 21 kPa, a temperature of 25°C, a basal growth medium originally free of carbonates, and an equilibrium pH of 6.0, no more than 10.5 mmol C per HRT period can be explained by dissolution of $CO_2$ in water.

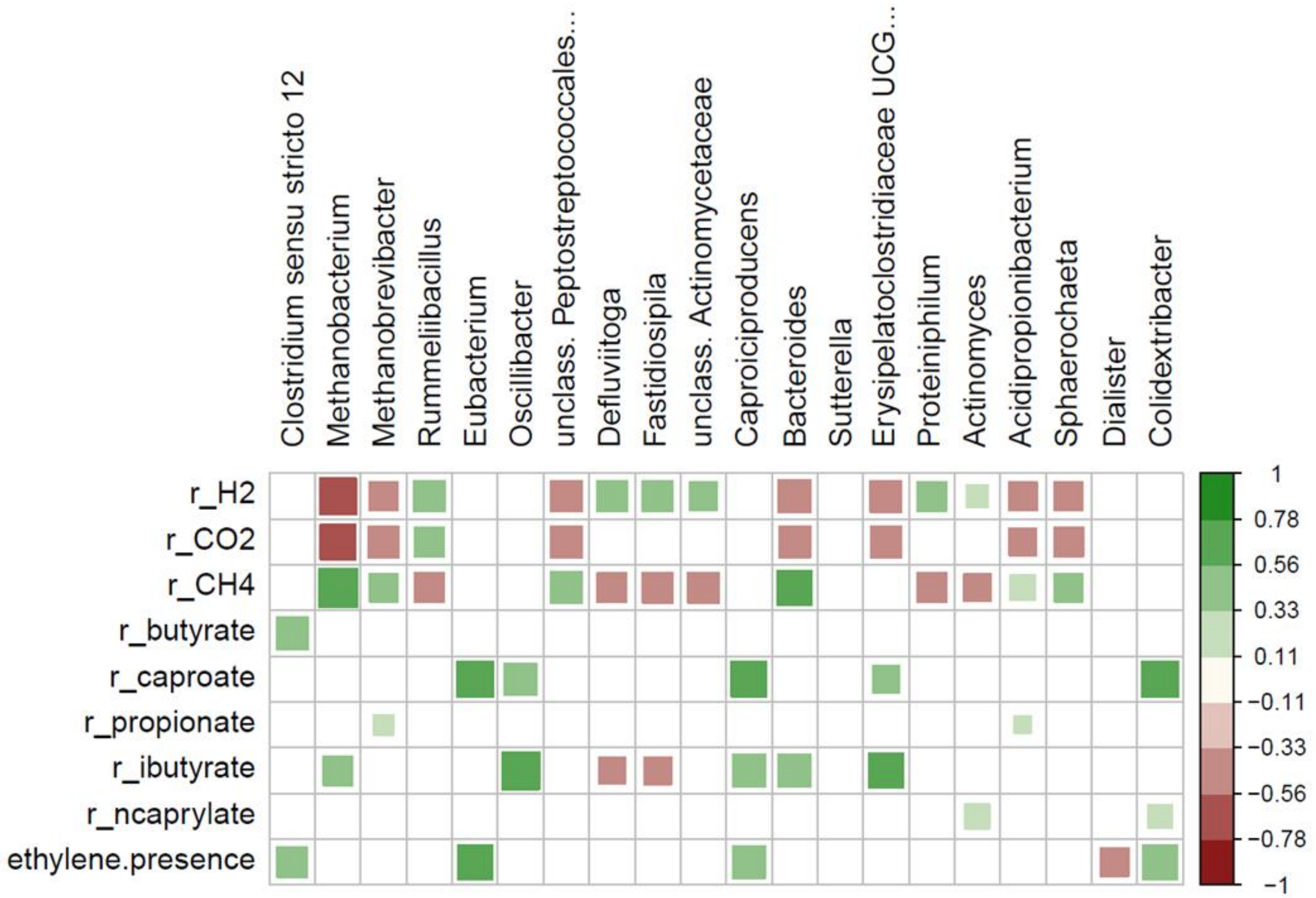


**Figure S9.** Correlation plot with the top 20 most abundant ASVs production rates of $H_2$, $CO_2$, $CH_4$, butyrate, caproate, propionate, *i*-butyrate, caprylate, and presence of ethylene. Spearman correlation is shown with $p<0.01$.

(A) (B)

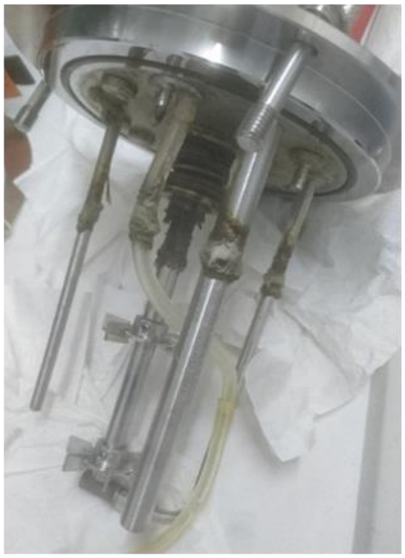

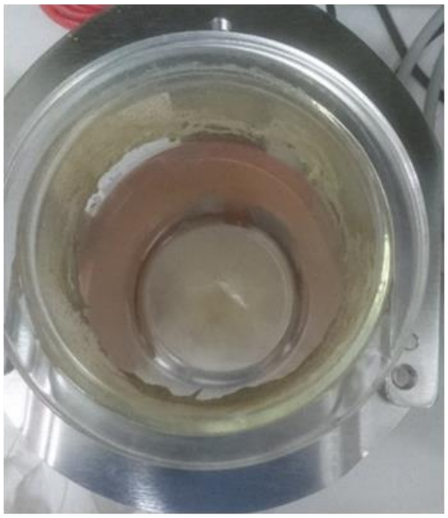

(C)

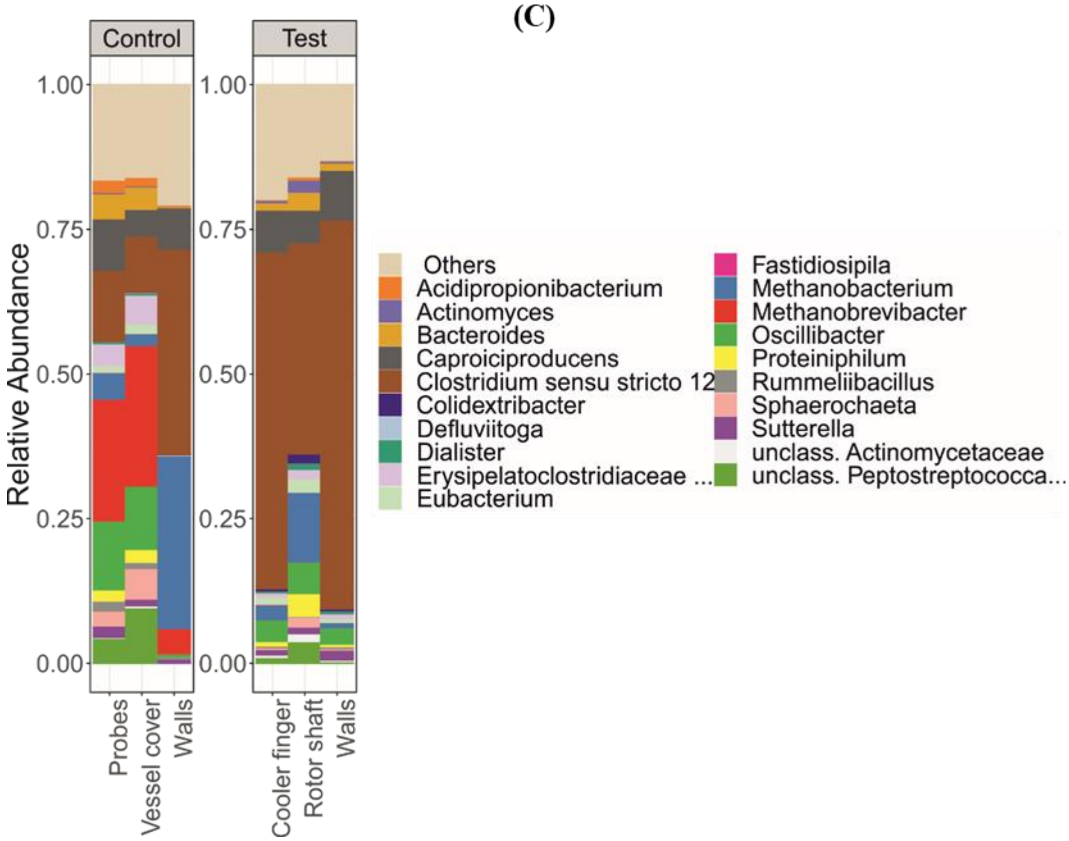


**Figure S10.** In biofilms attached to vessel cover, reactor probes, rotor shaft (A), and vessel walls (B) methanogens were detected regardless the use of ethylene (C). Methanogens were present in the planktonic community before ethylene addition but were washed out in the test reactor.

ASV SIMILARITIES TO SPECIES LEVEL. ASVs assigned to *Acidipropionibacterium* were related to *Acidipropionibacterium microaerophilum* (98.3% similarity of the V3-V4 regions of the 16S rRNA gene). Within the *Clostridium* sensu stricto 12 genus, ASVs were related to four different species. The most abundant clostridial species in both reactors were *C. tyrobutyricum* (99.8% similarity) followed by *C. luticellarii* (>99% similarity), a relative of *C. algifaecis* (>96.8% similarity), and *C. autoethanogenum* (100% similarity, with ambiguity to other species with identical V3-V4 16S rRNA gene: *C. ljungdahlii*, *C. ragsdalei*, and *C. coskatii*). The *Eubacterium* ASV had 98.3% similarity to *E. limosum* and the *Colidextribacter* ASV had 97.5% similarity to *C. massiliensis*. *Caproiciproducens galactitolivorans* was the most similar species to the *Caproiciproducens* ASV present in the system (94.1% similarity).

6. Angenent, L. T.; Richter, H.; Buckel, W.; Spirito, C. M.; Steinbusch, K. J.; Plugge, C. M.; Strik, D. P.; Grootscholten, T. I.; Buisman, C. J.; Hamelers, H. V., Chain Elongation with Reactor

Microbiomes: Open-Culture Biotechnology To Produce Biochemicals. *Environ Sci Technol* **2016,** *50* (6), 2796-810.
